# Citizen science, cells and CNNs – deep learning for automatic segmentation of the nuclear envelope in electron microscopy data, trained with volunteer segmentations

**DOI:** 10.1101/2020.07.28.223024

**Authors:** Helen Spiers, Harry Songhurst, Luke Nightingale, Joost de Folter, Roger Hutchings, Christopher J Peddie, Anne Weston, Amy Strange, Steve Hindmarsh, Chris Lintott, Lucy M Collinson, Martin L Jones

## Abstract

Advancements in volume electron microscopy mean it is now possible to generate thousands of serial images at nanometre resolution overnight, yet the gold standard approach for data analysis remains manual segmentation by an expert microscopist, resulting in a critical research bottleneck. Although some machine learning approaches exist in this domain, we remain far from realising the aspiration of a highly accurate, yet generic, automated analysis approach, with a major obstacle being lack of sufficient high-quality ground-truth data. To address this, we developed a novel citizen science project, Etch a Cell, to enable volunteers to manually segment the nuclear envelope of HeLa cells imaged with Serial Blockface SEM. We present our approach for aggregating multiple volunteer annotations to generate a high quality consensus segmentation, and demonstrate that data produced exclusively by volunteers can be used to train a highly accurate machine learning algorithm for automatic segmentation of the nuclear envelope, which we share here, in addition to our archived benchmark data.

## 1 Main

Until recently the study of cell morphology with electron microscopy (EM) was often restricted to qualitative illustration, as technological limitations prevented quantitative analysis of samples in three dimensions. Development of novel volume EM methodologies, including Serial Blockface Scanning Electron Microscopy (SBF SEM) [1] and Focused Ion Beam SEM (FIB SEM) [2], have enabled automated acquisition of images through greater depths at high resolution [3], with one microscope able to generate hundreds of gigabytes of aligned serial images per day.

However, our ability to analyse this data has not seen comparable advancement; segmentation of EM images remains a difficult and time-consuming manual process. Hence, to fully realize the analytical potential of EM, there is a great need to develop fast, generalizable and accurate analysis solutions. Although some EM image analysis can be automated through application of methods such as machine learning [4, 5, 6, 7, 8], these advances have mainly benefited specific domains such as connectomics [9, 10], where the segmentation problem is focused on tracing neurons and synapses in serial images from brain and nerves. This focus has generated a large amount of ‘ground truth’ data which has been successfully used in deep learning to generate algorithms to automate the task.

The same cannot be said of cell biology, where the segmentation challenge is more diverse, encompassing common organelles such as the nucleus, nuclear envelope (NE), mitochondria, endoplasmic reticulum and endosomes, as well as rare or transient organelles such as autophagosomes, secretory granules and phase-separated entities. As in connectomics, the production of ground truth segmentations has, to date, relied on the effort of the expert EM community. At present rates of data generation, this community alone is unable to generate sufficient ground truth segmentation data, representative of the appearance of the full range of organelles in different experimental conditions and biological model systems. To enable data analysis at a scale beyond the capacity of the research community, we engaged the help of a global community of willing volunteers through a novel online citizen science project, ‘Etch a Cell’ (www.zooniverse.org/projects/hspiers/etch-a-cell), which asked members of the public to manually segment the NE, which was targeted for volunteer segmentation as it is the most easily identifiable subcellular structure for which reliable automatic segmentation was not widely available.

The NE is a double lipid bilayer found in most eukaryotic cells where it surrounds the nucleoplasm and encloses the genetic material of the cell. Alterations in the structure of the NE have been associated with disease [11] including cancer [12, 13] and nuclear laminopathies [14]. However, despite the clear critical role of the nuclear envelope in cell function, the nanoscale three dimensional structure of this organelle has been poorly understood to date. In addition to its biological importance, segmentation of the NE is often a critical first step in the segmentation of a cell, as this structure provides important context to the three dimensional spatial distribution of other organelles.

Here, we present our method for establishing a high-quality consensus segmentation from multiple volunteer annotations on the same image. We demonstrate that exclusively volunteer produced data can be used to train a machine learning model for highly accurate automatic segmentation of the NE. Finally, we present a novel multi-axis modification of our machine learning algorithm that resulted in a marked improvement in model performance. We share all benchmark data and algorithms produced for the use of the wider research community.

## 2 Results

### 2.1 Etch a Cell: an online citizen science project for nuclear envelope segmentation

An online citizen science project, ‘Etch a Cell’ (EAC), was developed to enable large-scale segmentation of the NE in volume EM data through public engagement. Although online citizen science has been previously applied in similar contexts [15, 16], to our knowledge, this is the first application of non-expert, volunteer effort for the segmentation of organelles in EM data. To maximise the potential utility of the data produced for the research community, the commonly used HeLa cell line [17] was selected for analysis. A benchmark serial image data set was generated at 10nm pixel resolution with SBF SEM (Fig. 1, Supplementary Movie. 1), and n = 18 cells selected from this volume for volunteer segmentation (Supplementary Table. 1 details the unique Cell ID assigned to each ROI and provides further descriptive information), resulting in a total of n = 4241 slices for project inclusion after volume pre-processing (Supplementary Fig. 1A-D, and Methods). Raw data have been made available via the EMPIAR repository (deposition ID: 137, accession code: EMPIAR-10094, www.ebi.ac.uk/pdbe/emdb/empiar/entry/10094).

**Figure 1:**
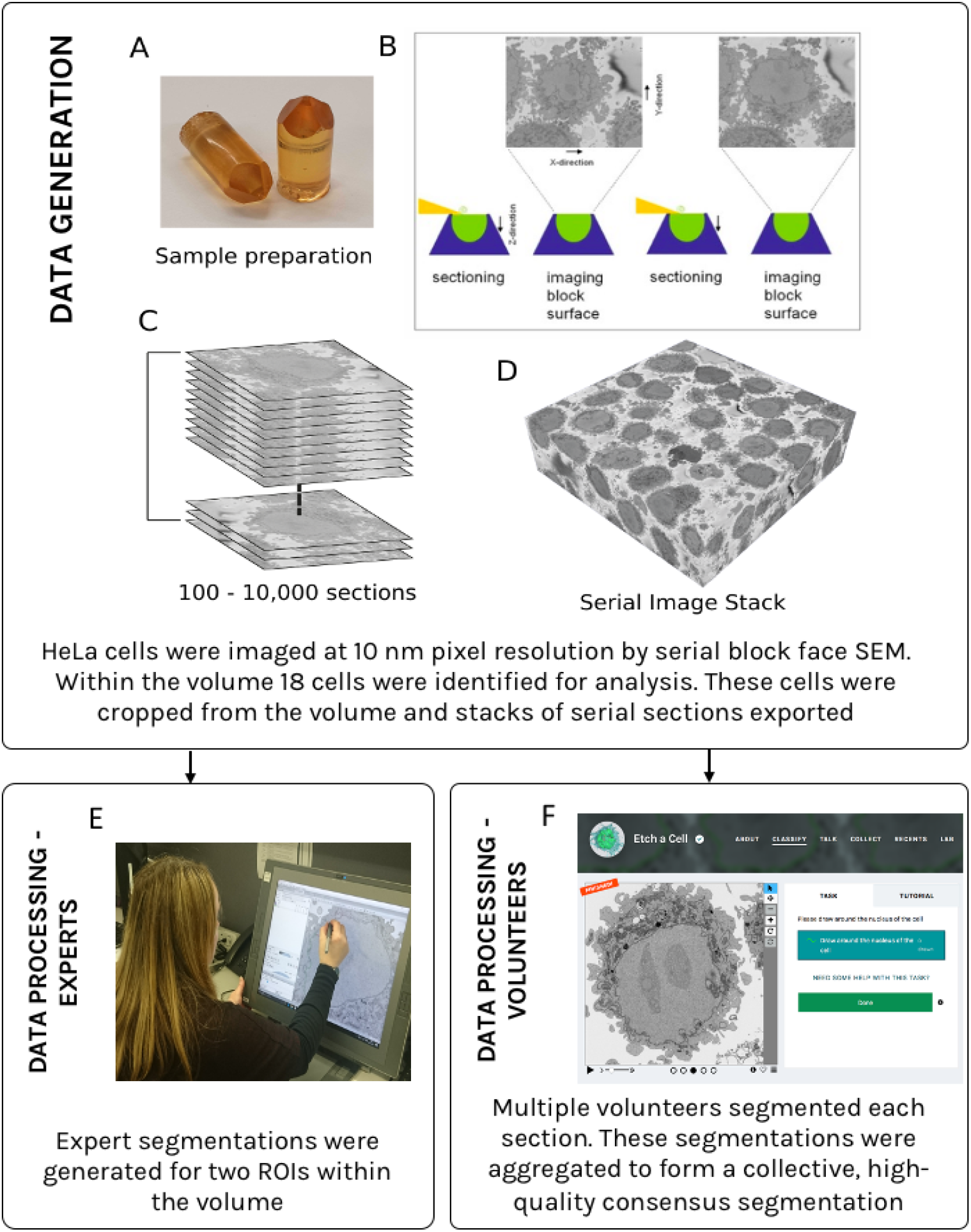
Workflow for the acquisition and segmentation of serial EM images from benchmark samples. In this study we imaged resin-embedded HeLa cells at 10nm pixel resolution (A) using SBF SEM (B). This produced an image stack (C) of 518 sections (50nm thickness, 8192 × 8192 pixels, Supplementary Movie. 1) which were used to construct a 3D volume (D). ROIs from within this volume were segmented by both experts (E) and volunteers (F). Supplementary Table. 1 provides further information about the individual ROIs within the volume.

### 2.2 Development and deployment of Etch a Cell

Following an iterative design process lasting approximately six months (from October 2016 to the beginning of April 2017), EAC was launched on 6th April 2017 as a publicly available project on the Zooniverse online citizen science platform (www.zooniverse.org). Volunteers could contribute segmentations through visiting an online classification interface (Supplementary Fig. 1E), where they were presented with a cell slice at random. A detailed tutorial provided task instructions (Supplementary Fig. 2), describing how to segment the NE with the novel ‘Freehand Drawing Tool’ and how to resolve segmentation ambiguities through viewing the two neighboring slices either side of the target slice (Supplementary Fig. 1C-E, and Methods). Each slice was independently segmented by multiple volunteers with the number of classifications required set at n = 30 (the ‘retirement limit’). Each individual segmentation could consist of an arbitrary number of lines, which were recorded as an array of x,y pairs (Methods, Supplementary Fig. 1F).

### 2.3 An overview of volunteer interaction with Etch a Cell

In total, n = 104612 classifications were submitted by volunteers before the EAC workflow was deactivated on 1st August 2019. As classifications could be made by unregistered volunteers, it was not possible to establish precisely how many individuals contributed, however, classifications submitted by logged in users were associated with n = 4749 user ids and n = 9444 IP addresses, indicating between 5000 - 10000 individuals contributed. As is often observed for online citizen science projects [18], a large number of classifications were received shortly after project launch (Supplementary Fig. 3A), and the number of classifications submitted by each volunteer varied greatly (from 1 to 5451), indicating a broad range of engagement levels amongst the community contributing to this project (Supplementary Fig. 3B). Examining the Lorenz curve for the distribution of volunteer classification contributions to the project (Supplementary Fig. 3C) and the corresponding Gini coefficient (0.83), indicated that a small number of volunteers contributed a large number of classifications, as is commonly observed in citizen science projects [18]. However, it should be noted that a significant proportion of the classifications submitted to the project were made by individuals only submitting a small number, reiterating the importance of all individual contributions.

### 2.4 Forming consensus from multiple segmentations - aggregating volunteer annotations

To generate sufficiently high quality data for downstream analyses, each individual slice within EAC was presented to multiple volunteers for segmentation. As expected, most volunteer segmentations were distributed on and around the NE (Fig. 2A-C and Supplementary Movie. 2), however, distinct classes of segmentation error were observed, including ‘graffiti’ (Fig. 2D, possibly produced as an inadvertent consequence of well-intended classroom based engagement using this project), ‘false positive segmentation’ in which non-NE pixels are segmented (Fig. 2E) and ‘false negative segmentation’ where NE pixels are missed (Fig. 2F). Of these error classes, the graffiti class was comparatively rare (Supplementary Movie. 2).

**Figure 2:**
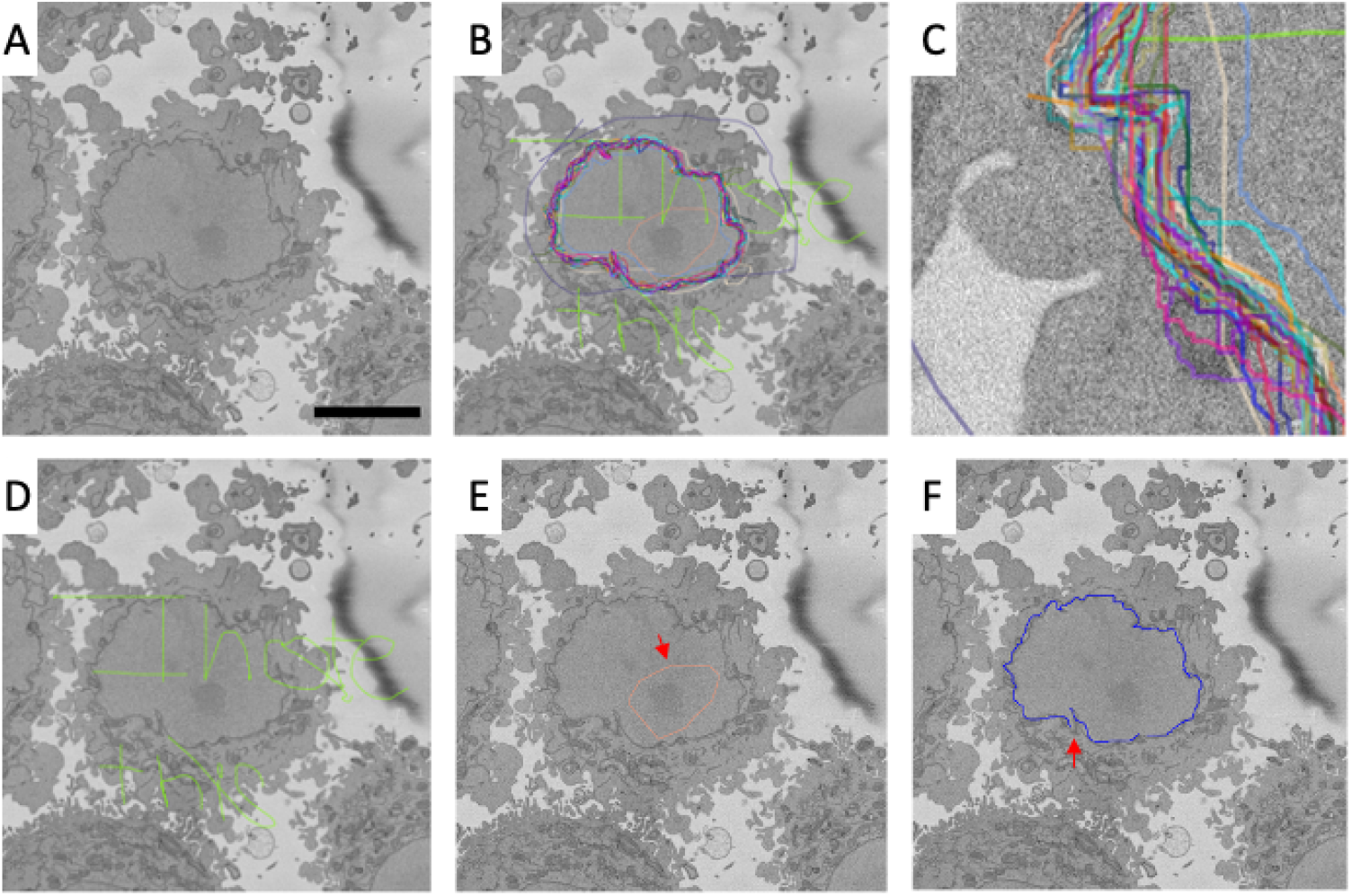
Multiple segmentations were contributed by volunteers for each image in the EAC project. Each cell slice (e.g. A) was segmented by multiple volunteers (contributions from different volunteers shown in different colours (B) and zoomed in panel (C)). The volunteer annotations vary in quality, with identifiable classes of error including; graffiti (D), ‘false positive segmentation’ where non-NE is segmented, e.g. a region within the NE, indicated with a red arrow (E) and ‘false negative segmentation’ where NE pixels have been missed, indicated with a red arrow (F) (Supplementary Movie. 2). For downstream analyses it is necessary to aggregate the multiple volunteer segmentations to establish a final ‘consensus’ NE segmentation for each slice. Panels are produced from slice number 70 from C001 (ROI 1656-6756-329) and 5 micron scale bar is shown on panel A.

To remove outlying data and establish a ‘consensus’ segmentation for each slice, it was necessary to aggregate the multiple volunteer annotations. As the Freehand Drawing Tool was developed specifically for EAC, it was necessary to develop a novel aggregation approach. Due to the presence of noise, erroneous segmentations, and an unknown, variable number of line segments within the data, this was not trivial, hence, multiple novel aggregation approaches were developed and explored. Of these, the ‘Contour Regression by Interior Averages’ (CRIA) algorithm was selected for our analytical pipeline, as it had a number of advantages compared to other approaches as shall be outlined.

Briefly, the CRIA algorithm procedure involved the following steps: first, closed loops were formed from each individual volunteer segmentation (Fig. 3A-C), which could consist of multiple separate lines (Supplementary Fig. 1F). The closed loops were produced through connecting separate lines after ordering them by minimizing distances. Next, interior areas were generated from the closed loops (Fig. 3B). The interior areas were overlaid to generate a height map, with the ‘height’ reflecting the level of agreement between the separate volunteer segmentations for a single slice (Fig. 3D). The consensus segmentation was determined through taking a mean ‘height’ level, hence the resulting, ‘final’, segmentation surrounds all the interior areas where half or more of the volunteer segmentations were in agreement (Fig. 3E-F). This procedure was used to aggregate all volunteer segmentations (Fig. 3G, Supplementary Movie. 3). The resulting aggregated data has been made available at www.ebi.ac.uk/biostudies/files/S-BSST448/Aggregations and the CRIA code is available at www.github.com/FrancisCrickInstitute/Etch-a-Cell-Nuclear-Envelope.

**Figure 3:**
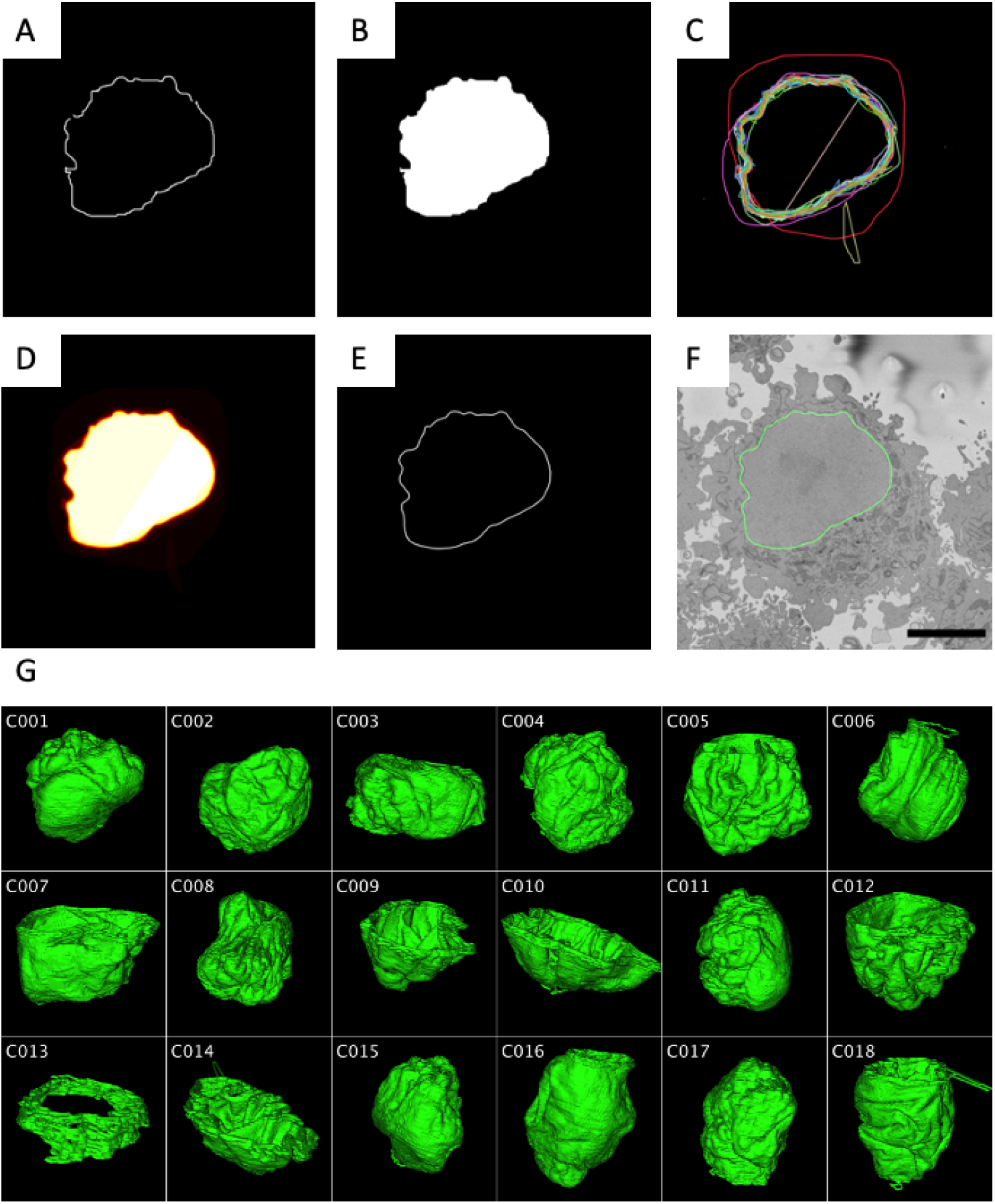
The Contour Regression by Interior Averages (CRIA) algorithm was developed for the aggregation of multiple volunteer segmentations. In this algorithm, each individual volunteer segmentation (A) was converted into a closed loop (B). This procedure was performed for all the segmentations associated with each slice of the ROI, as can be seen stacked in (C). The closed loops were converted to interior areas and stacked (D). A final, consensus segmentation was determined as the outline of all interior areas where half or more of the volunteer segmentations were in agreement (E). This generated a high-quality, volunteer-produced segmentation (5 micron scale bar) (F). We show here the annotations and aggregation for slice number 150 from C001 (ROI 1656-6756-329). This process was applied to each slice of all n = 18 volunteer-segmented ROIs, allowing generation of a 3D reconstruction of each ROI (G) (Supplementary Movie. 3).

In addition to producing high quality consensus segmentations (Fig. 3F-G) that showed high visual similarity to expert data (Fig. 4A, B, D, E, G, H, Supplementary Movie. 4 and Supplementary Movie. 5) this aggregation procedure had a number of notable benefits. Importantly, the CRIA algorithm made use of all volunteer annotations in producing the final consensus segmentation, therefore no volunteer effort went unused. In contrast to other methods explored, the CRIA algorithm is fully automated and required no expert involvement, such as manual selection of high quality segmentations to seed the algorithm. This is a critical advantage for a number of reasons; avoiding a requirement for manual intervention minimizes the possibility of perturbing the final, consensus, segmentation through introducing subjectivity and bias associated with a single individual. Further, eliminating the need for expert assessment removes a significant analytical bottleneck. Finally, and perhaps most notably, this approach has shown high quality segmentations can be generated solely through collective non-expert effort.

**Figure 4:**
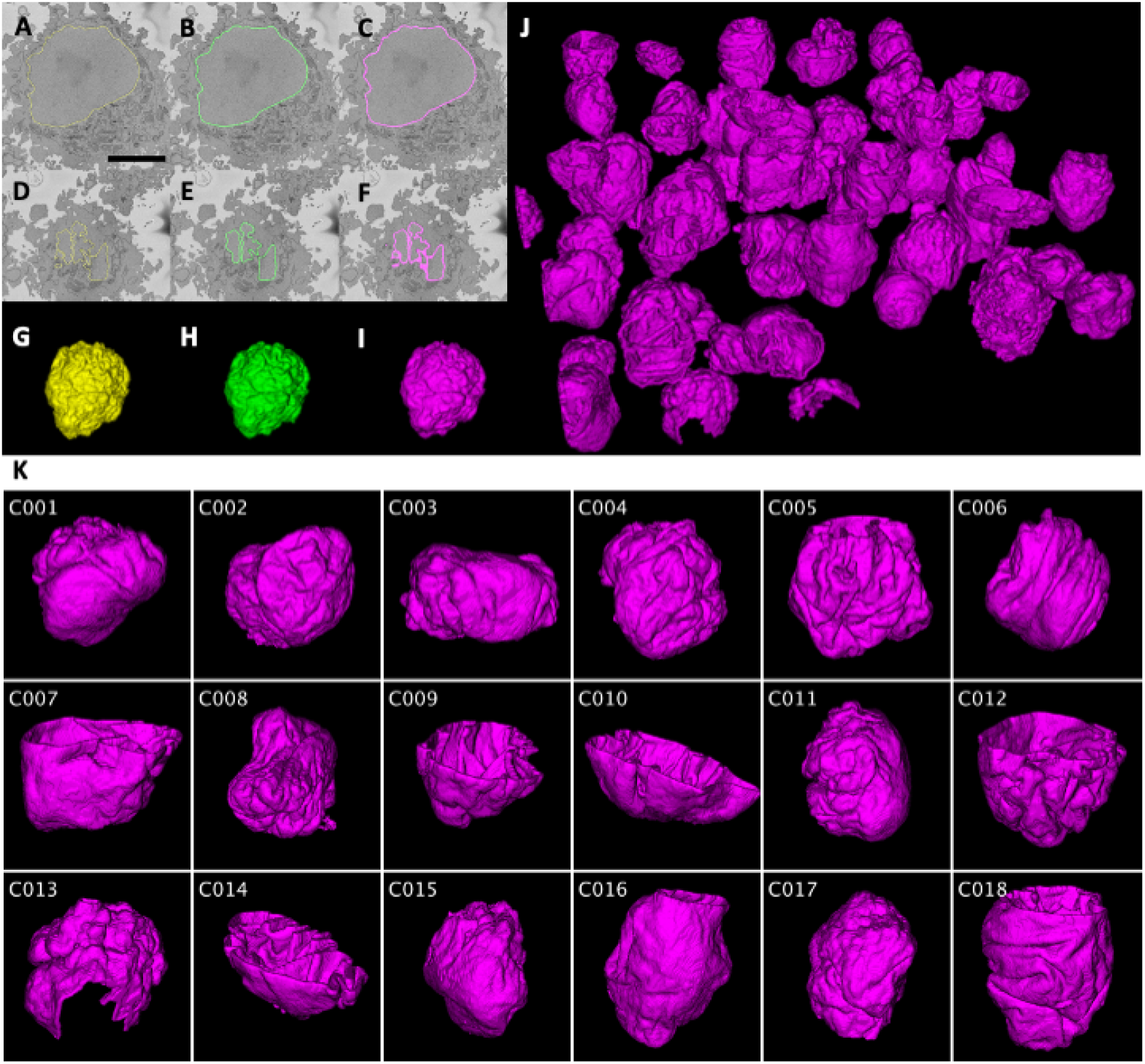
Consensus volunteer and machine-predicted NE segmentations are high quality. Visual inspection reveals a high similarity between expert (A), aggregated volunteer (B), and machine predicted (C) segmentations (shown for slice number 150 from C001 (ROI 1656-6756-329)), and a high degree of overlap of these segmentations with the NE. Segmentations from slices found at the top and bottom of the volume (D, E, F) showed greater segmentation variability due to the presence of NE islands and membrane parallel to the cutting plane, which make these regions more challenging to segment (shown for slice number 40 from C001 (ROI 1656-6756-329)). 5 micron scale bar is shown on panel A. Despite this, 3D reconstruction of nuclei revealed a high similarity between expert (G) volunteer (H) and machine (I) segmented nuclei (shown for C001 (ROI 1656-6756-329)) (Supplementary Movie. 4 and Supplementary Movie. 5). Automatic NE segmentation using our trained model applied to our full data volume captured many nuclei which had not previously been segmented through expert or volunteer effort (J) (Supplementary Movie. 9), and the n = 18 nuclei previously segmented by volunteers (K) (Supplementary Movie. 10). Machine-predicted segmentations were produced with TAP.

### 2.5 Machine learning for NE segmentation

Aggregated volunteer NE segmentations were used to train a U-Net CNN architecture [19, 20] for automatic segmentation of the NE in SBF SEM data. Model performance was assessed through presenting the model with two previously unseen ROIs, and comparing the resulting predicted NE segmentations to ‘ground truth’ (Supplementary Table. 1 and Methods provide further information about ROIs used for model training, validation and testing).

Two complementary forms of ‘ground truth’ data were available; expert generated segmentations (available at www.ebi.ac.uk/biostudies/files/S-BSST448/Expert) and aggregated volunteer segmentations (Methods, Supplementary Table. 1 and Fig. 4) providing a means to test two facets of model performance. In comparing the prediction to the aggregated volunteer data for each ROI, we were able to establish how well the model had learnt to perform the task of NE segmentation from the training data provided, which consisted of exclusively volunteer produced segmentations. Comparing model performance to expert data enabled assessment of how well the model (hence, indirectly, the volunteers) performed this task in comparison to experts.

The model performed well when compared to aggregated volunteer data. The Average Hausdorff Distance (AHD) between the predicted segmentation and the aggregated volunteer segmentation was 1.638 pixels (corresponding to a distance of 16.377nm) for C001, and 1.767 pixels (17.675nm) for C006. The F-measure, recall and precision of the model were 0.700, 0.792, 0.628 respectively for C001 and 0.687, 0.767, 0.621 for C006 (Table. 1). Although these metrics may initially seem poor in comparison to similar previous work [21, 22], it should be emphasised that we are examining the overlap between lines (the NE) rather than areas (the nucleus). Hence, for easier comparison of our model with previously reported metrics, we also provide the F-measure for the nucleus area for a single slice within each ROI. As anticipated, this metric shows a much higher model performance of 0.995 (C001) and 0.991 (C006).

**Table 1:**
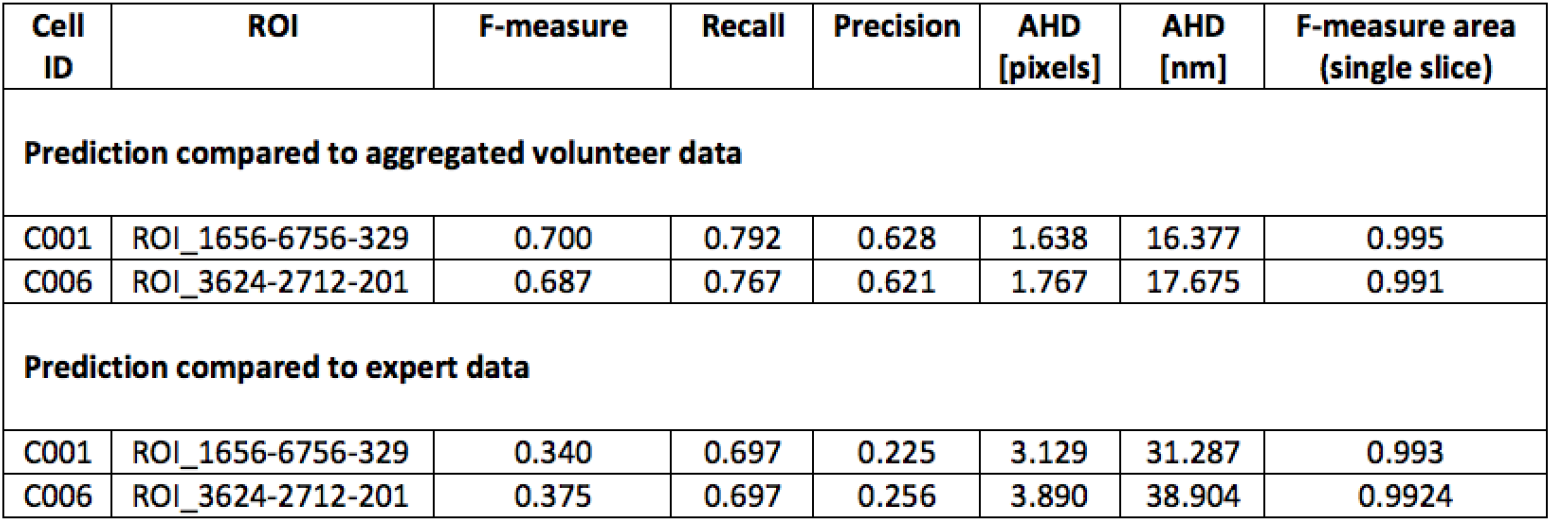
CNN performance metrics. Model performance was assessed by comparing the predicted NE segmentation to ground truth for two ROIs (ROI 1656-6756-329 (Cell ID = C001) and ROI 3624-2712-201 (Cell ID = C006)). We report multiple metrics of model performance (Methods) against two complementary modes of ‘ground truth’ data available (aggregated volunteer data and expert-produced segmentations).

| Cell ID | ROI | F-measure | Recall | Precision | AHD [pixels] | AHD [nm] | F-measure area (single slice) |
| --- | --- | --- | --- | --- | --- | --- | --- |
| **Prediction compared to aggregated volunteer data** |  |  |  |  |  |  |  |
| C001 | ROI_1656-6756-329 | 0.700 | 0.792 | 0.628 | 1.638 | 16.377 | 0.995 |
| C006 | ROI_3624-2712-201 | 0.687 | 0.767 | 0.621 | 1.767 | 17.675 | 0.991 |
| **Prediction compared to expert data** |  |  |  |  |  |  |  |
| C001 | ROI_1656-6756-329 | 0.340 | 0.697 | 0.225 | 3.129 | 31.287 | 0.993 |
| C006 | ROI_3624-2712-201 | 0.375 | 0.697 | 0.256 | 3.890 | 38.904 | 0.9924 |

Reflecting differences in volunteer and expert segmentation skill, it was expected that we would see reduced model performance (trained exclusively using volunteer produced data), when comparing against expert-produced ‘ground truth’ data. We observed an AHD of 3.129 pixels (31.287nm) for C001 and 3.890 pixels (38.904nm) for C006 between the prediction and expert data. The F-measure, recall and precision were 0.340, 0.697, 0.225 respectively for C001, and 0.375, 0.697, 0.256 for C006 (Table. 1). Although most of these metrics indicate good model performance, the F-measure and precision warrant further explanation. These measures are particularly poor in the case of comparing the model to the expert data due to an idiosyncrasy of the expert data. The width of the available expert data is narrower (30nm) compared to both the aggregated and predicted width of the NE (70nm), and because of this we see a degradation of the precision and F-measure metrics. This is because the model has assigned pixels as NE that do not correspond with pixels annotated by the expert, therefore, the false positives rate is seemingly inflated. Unfortunately, it was not feasible to amend our expert ground truth through either asking an expert to resegment the ROIs (this was not practical due to time constraints) nor was it recommendable to dilate the width of the expert segmentation (as this would introduce greater errors e.g. incorrectly assigning cytoplasm pixels as NE). Despite this, upon visual inspection, it was found that the model performance was arguably superior to the expert segmentation as more relevant pixels appeared to be assigned to the NE by the model (Fig. 4), which raises questions regarding the legitimacy of ‘ground truth’ data produced by a single expert, as shall be discussed later.

### 2.6 Improved model performance with Tri-Axis Prediction

Although the model performed well, expert visual inspection revealed some regions of undersegmentation (Supplementary Fig. 4A). These regions were not randomly distributed across the data, but were instead localized to sites at the top and bottom of the volume (the highest and lowest z slices, Supplementary Fig. 4B,C), presumably due to the higher degree of visual ambiguity in these regions caused by the presence of a greater number of NE islands and the membrane being oriented parallel to the SBF SEM imaging plane. To improve the automated segmentation, we sought to leverage additional information available in the volume. The data examined here were downscaled in the xy plane to 50nm to be isotropic, therefore, it was possible to transpose the stack and run the model on each axis (Supplementary Fig. 4D-F). This resulted in three orthogonal NE predictions which were recombined to form a final segmentation, with pixels assigned to NE in all three predictions accepted (Supplementary Fig. 4G, Supplementary Movie. 6 and Supplementary Movie. 7) and over-segmented pixels removed using a connected components analysis (Supplementary Fig. 4H). Visual inspection revealed a significantly improved segmentation (Supplementary Fig. 4I), however, it was not possible to quantify this improvement due to a lack of appropriate ground truth data.

The Tri-Axis Prediction (TAP) approach was applied to the entire volume (Fig. 4, Supplementary Movie. 1, Supplementary Movie. 8 and Supplementary Movie. 9), and took a total of 48 minutes to produce NE predictions for all nuclei within the volume (Fig. 4J), including the n = 18 nuclei already segmented with volunteer effort (Fig. 4K and Supplementary Movie. 10). TAP predictions have been made available at www.ebi.ac.uk/biostudies/files/S-BSST448/Predictions. Serendipitously, a cell within the volume was undergoing mitosis, allowing us to observe that our model performed well in this challenging context in which the NE had partially broken down, despite not having been exposed to training data of this type (Fig. 5A and Supplementary Movie. 11). This is in contrast to some other approaches for NE identification which rely on the presence of a clear boundary, such as flood or marker based watershed methods [23]. The ability of the algorithm to segment disassembled mitotic NE is particularly surprising given the NE effectively regresses to become ER during mammalian cell division. Further analysis of the features identified by the model may be useful in defining the transition of the NE to the ER and back during mitosis. TAP was also applied to an alternative region from the same resin-embedded sample imaged at higher resolution (5nm) on the same microscope (which also contained both mitotic and interphase cells, Fig. 5B and Supplementary Movie. 12), and to a HeLa cell from the same sample imaged by an alternative volume EM methodology (FIB SEM) (Fig. 5C and Supplementary Movie. 13). Visual inspection of these data sets showed good model performance indicating the model is generalisable to novel contexts, however, it should be acknowledged that some erroneous over-segmented pixels can be observed, particularly in the peripheral ER and edges of lipid droplets bordering the nuclear region in mitotic cells (Fig. 5B and Supplementary Movie. 12), indicating there is scope for future improvement.

**Figure 5:**
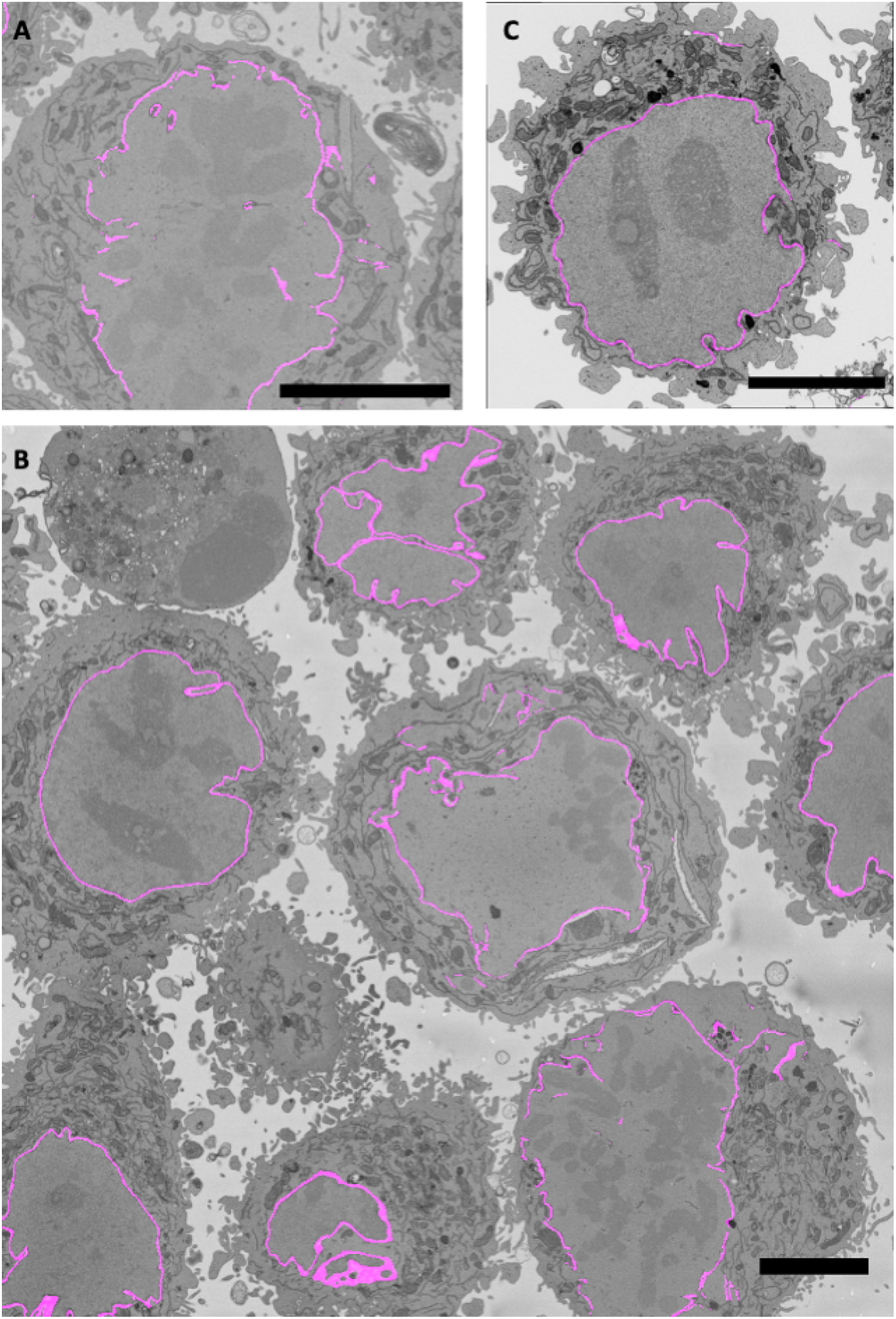
Model shows high NE segmentation performance in novel contexts. Applying TAP to the full data volume allowed us to observe that the model performed well when segmenting the partially broken down NE of a mitotic cell (A) (Supplementary Movie. 11). TAP also showed good performance when applied to the same resin-embedded sample (which also contained both mitotic and interphase cells) imaged at higher resolution (5nm) on the same microscope (B) (Supplementary Movie. 12) and when applied to a HeLa cell from the same sample imaged by an alternative volume EM methodology, FIB SEM (C) (Supplementary Movie. 13). All scale bars are 5 micron.

Finally, considering the overarching objective of this research was to facilitate faster analysis of volume EM data, we calculated the time benefit of our predictive model. For an expert to manually segment a single ROI would have taken approximately 30 hours [24]. Prediction took approximately 1 minute per image stack for a single ROI, therefore, our model prediction represents 1800 times faster data analysis.

## 3 Discussion

We show here that volunteer effort through online citizen science can be effectively applied to the task of manual segmentation of organelles in electron micrographs, enabling data analysis at a scale not achievable by experts alone. We demonstrate the data produced is of sufficient quality for task automation through training a CNN capable of segmenting the NE at high accuracy. Although prior work has shown crowdsourced volunteer effort can be productively applied to comparable tasks, such as the marking of single particles from cryo-EM micrographs to generate 3D protein reconstructions [15], and to the marking of whole cells [16], to our knowledge this is the first study to demonstrate the ability of volunteers to effectively perform manual freehand segmentation of an organelle in volume EM data.

Such large-scale, systematic segmentation makes quantitative examination of organelle morphology feasible. This has the potential to drastically advance our understanding of NE morphology and function, in both normal and diseased states such as cancer [12, 13] and nuclear laminopathies [14]. Yet, even with the collaboration of a community of citizen scientists it will not be possible to segment data at a scale proportional to current data production rates, and this challenge will become greater with further technological advancement. Hence, we sought to automate NE segmentation using volunteer produced segmentations as training data for a CNN [19], resulting in a model able to segment the NE to a high standard in a matter of minutes, rather than the hours, days or weeks required for manual segmentation. Critically, our model was trained exclusively with volunteer segmentations and required no expert microscopist input or intervention.

Although our model performed surprisingly well when applied to data produced under different conditions, there was scope for improved performance. Hence, we remain far from our aspiration of a highly-accurate, yet broadly applicable approach for the automated analysis of microscopy data for a single organelle, let alone each feature of interest in every volume acquired. It is anticipated that future work in this arena may be accelerated through application of approaches including transfer learning [25] and multiclass predictive models [26]. However, unless significant advances are seen in unsupervised machine learning approaches, it is expected that an adequately trained generic model will still require a large quantity of ground truth annotations, covering each organelle from a wide variety of imaging conditions. Clearly, the lack of suitable training data remains a significant barrier to the development of a broad automated approach, however, as we have demonstrated here, large training data sets can be produced through volunteer engagement.

Many avenues exist for the extension of this research, including segmentation of other organelles, examination of data from other imaging modalities, and the analysis of data from further cell lines or tissue types. We anticipate that such future projects would be significantly expedited by the workflows and analyses established in this first iteration of EAC. However, consideration will be required to effectively apply the methods and tools presented here to novel contexts. A significant degree of variation exists in the difficulty of manual segmentation of different organelles under different conditions, we will therefore need to refine and modify novel projects. For example, in designing our second EAC project for the segmentation of mitochondria (‘Etch a Cell - Powerhouse Hunt’ www.zooniverse.org/projects/h-spiers/etch-a-cell-powerhouse-hunt) it was necessary to adjust the field of view presented to the volunteer to ensure a reasonable number of mitochondria of an appropriate size were presented for segmentation. For organelles that can only be unequivocally identified by functional markers (e.g. autophagosomes labelled with GFP) other study design adjustments will need to be explored such as presenting correlative light images in conjunction with EM images to guide segmentation.

Although challenging from a study design perspective, the possibility of designing a portfolio of projects of varying difficulty provides a rich opportunity to engage volunteers through serving a greater variety of skills and interests. Reassuringly, citizen scientists have proven capable of performing a growing array of challenging tasks, from identifying supernovae [27] to visually assessing the quality of brain registration in fMRI studies [28]. We have been astonished that it is possible to train non-experts to recognise and segment complex organelles in minutes with just an online tutorial. We therefore remain confident in the abilities of our volunteer community to successfully perform novel segmentation tasks.

The challenge of motivating increased engagement and high-quality contributions will become increasingly important as our repertoire of citizen science projects expands and diversifies. Manual segmentation is a challenging task requiring a large time investment. We must therefore continue to develop novel modes of engaging our community and work to reduce the effort required to segment each slice. Mechanisms to achieve reduced volunteer effort per slice may include ‘smart subject assignment’ [29, 30] – the intelligent passing of slices of appropriate difficulty to volunteers. Further, it may be possible to actively retire slices from the project once an acceptable segmentation quality has been achieved – this would enable volunteer effort to be reduced for ‘easier’ images, and to instead be applied to images of greater difficulty.

Incorporating ‘computers in the loop’ may provide additional mechanisms for reducing segmentation effort. Future pipelines may include presenting volunteers with predicted segmentations for correction, rather than full segmentation. Feedback loops between computer-prediction and crowd-correction could enable real-time model refinement, improve predictions and therefore progressively-reduce need for volunteer correction, resulting in greater project efficiency. Predictive models need not be fully optimised to be useful; if a model is not yet able to accurately segment its target organelle, it may still provide valuable information that could be fruitfully leveraged, for example, the anticipated number of a particular organelle class and their approximate location. This would provide a mechanism for assessing volunteer ability, segmentation quality and subject difficulty.

We have demonstrated that experts can be removed from the task of manual segmentation, however, researcher time remains necessary to generate the infrastructure supporting this effort and for the continued refinement of multiple aspects of the analytical pipelines underlying these studies. More critically, researcher effort continues to be needed to interpret and assess the quality of volunteer or machine produced segmentations. This is a particularly challenging component of this work, as the required quality of a ‘final segmentation’ is often intimately linked to the research question being addressed. Related to the ubiquitous challenge of finalising segmentations and establishing ‘truth’ in this domain: it can be difficult to definitively assign pixels of noisy and nuanced micrographs to different regions, and much inter- (and intra-) expert variation can exist. The potential for pixel-assignment disagreement raises an interesting possibility regarding additional value of collectively producing segmentations; when multiple individuals annotate each slice, rather than a single expert, it is possible to produce a level of confidence that each pixel belongs to a certain region, rather than simply a binary designation. Such a segmentation-confidence map may be more reflective of the reality of cell morphology, where a subset of pixels may not definitely belong to a particular region. This may provide insight, with regions of variable confidence being of possible biological relevance, for example, we may expect nuclear pores in the NE to be less confidently designated as this organelle.

Future collaboration of the crowd and computing is poised to enable, for the first time, the large-scale, generalizable, yet accurate, quantification of multiple subcellular structures across many data modalities at the nanoscale.

## 4 Online Methods

### 4.1 Cell model

HeLa cells were grown in culture then fixed in 2.5% glutaraldehyde and 4% formaldehyde in 0.1 M phosphate buffer (pH 7.4), and embedded as a pellet in Durcupan resin following the method of the National Centre for Microscopy and Imaging Research [31].

### 4.2 Data acquisition

Serial blockface scanning electron microscopy (SBF SEM) data was collected using a 3View2XP (Gatan, Pleasanton, CA) attached to a Sigma VP SEM (Zeiss, Cambridge). Small portions of the cell pellets were mounted on pins using conductive epoxy resin (Circuitworks CW2400), trimmed to form an approximately 400 × 400 × 150 *μ*m pillar, and coated with a 2nm layer of platinum. Images were acquired at 8192 × 8192 pixels using a dwell time of 6 *μ*s (10nm reported pixel size, horizontal frame width of 81.99 *μ*m) and 50nm slice thickness. The SEM was operated at a chamber pressure of 10 pascals, with high current mode inactive. A 20 micron aperture was used, with an accelerating voltage of 3 kV.

Raw data consisted of a total of 518 images acquired sequentially, representing a depth of 25.9*μ*m and total volume of 174135*μ*m^3^ (10nm dataset, Fig. 1 and Supplementary Movie 1). One image was excluded from analysis due to a technical fault resulting in loss of the cut material and the production of a blank image. To enable further analyses and benchmarking, raw data have been made available via the EMPIAR repository (deposition ID: 137, accession code: EMPIAR-10094, www.ebi.ac.uk/pdbe/emdb/empiar/entry/10094). Digital Micrograph (DM4) files were read into Fiji with the Bio-Formats library [32] and subsequently saved as TIFFs.

### 4.3 Expert production of ground truth data

Ground truth segmentations for two ROIs were obtained by manual annotation in the Amira software package [33]. The two cells segmented by expert effort were Cell ID = C001 (ROI 1656-6756-329) and Cell ID = C006 (ROI 3624-2712-201) (Supplementary Table. 1). Expert segmentations have been made available at www.ebi.ac.uk/biostudies/files/S-BSST448/Expert.

### 4.4 Volume pre-processing to produce images for online citizen science

Approximately 40 cells were visualized within the full volume (Supplementary Movie 1). Of these, cells with nuclei not intersecting the edge of the field of view in any slice were manually selected for study inclusion. This ensured each 2D slice presented to volunteers contained a complete NE, simplifying the labelling of this structure and reducing the likelihood of misidentification. A total of n = 18 appropriate cell volumes were selected using this criterion. Each selected cell was cropped from the full volume (Supplementary Fig. 1A-C) and exported as a sequence of TIFF images using the following procedure within Fiji software; a 3D region of interest was created around each selected cell, with a size of 2000 × 2000 pixels in x,y. The number of slices (synonymous with ‘subjects’ in Zooniverse terminology) per cell ranged from 150 to 300 due to inherent differences in cell size and variability in cell completeness across the z-axis in the data volume (12 of the 18 cells weren’t complete in the z-axis, Supplementary Table. 1 and Supplementary Movie 1). To ensure reasonable web browser download times, the raw images for each ROI were downscaled to 1000 × 1000 pixels in x,y, and converted from Digital Micrograph DM4 format to JPEG with quality of 90% to reach an image size of approximately 600 KB using the Fiji software package [34] with the Bio-Formats tool [32].

### 4.5 Development of Etch a Cell with the Zooniverse Project Builder

Cell images were presented to volunteers for NE segmentation via an online citizen science project, ‘Etch a Cell’. This project was designed and deployed on the Zooniverse platform (www.zooniverse.org) using the Project Builder (www.zooniverse.org/lab). The Zooniverse Project Builder is a free, web-browser based toolkit that provides the core infrastructure necessary for designing, building and implementing all components of an online citizen science project, including the project workflow and supporting materials such as the project tutorial (Supplementary Fig. 2). Project development took approximately six months, during this time the project workflow was designed and all supporting materials produced, including an in-depth tutorial to comprehensively explain the NE segmentation task (Supplementary Fig. 2). Prior to launch, the project was refined through a multi-step review process involving thorough assessment of the project by both Zooniverse volunteers and the Zooniverse research team. For key Zooniverse terms please refer to help.zooniverse.org/getting-started/glossary.

### 4.6 Etch a Cell workflow design

The pre-processed slices were uploaded to EAC for volunteer segmentation using a python script (www.github.com/FrancisCrickInstitute/Etch-a-Cell-Nuclear-Envelope) that interfaced with the Zooniverse Panoptes API (www.github.com/zooniverse/Panoptes). Slices were embedded within the project ‘workflow’. The ‘workflow’ of a Zooniverse citizen science project refers to the series of tasks a volunteer is asked to complete when presented with data in the project’s classification interface.

In the EAC project workflow, upon being presented with one of the uploaded cell slices at random, volunteers were asked to perform the task of segmenting the NE using a Freehand Drawing Tool applied directly to the image in a web browser (Supplementary Fig. 1E). Upon submission of the classification, the individual lines drawn by the volunteers were recorded as arrays of x,y pairs defining a line path (Supplementary Fig. 1F).

To support volunteers in the task of NE segmentation, a detailed project tutorial was provided on the classification interface (Supplementary Fig. 2). To enable more accurate annotation, pan and Zoom functionality was enabled. Further, to provide the volunteer with a limited amount of three-dimensional context to help resolve segmentation ambiguities, the image to be segmented was presented as the central image within a ‘flipbook’ of five images, with the two neighbouring images on either side corresponding to the +/− 250 and 500nm planes in the z-dimension (Supplementary Fig. 1D, E)

To produce data of sufficient quality for downstream analyses, each individual image was presented to multiple volunteers to enable the generation of a ‘consensus’ from the aggregation of multiple annotations. The minimum number of required annotations is denoted the ‘retirement limit’, and was set at n = 30 for this project. Therefore, each individual image received at least 30 volunteer segmentations. A small subset of images received more than this, as a small number of images continue to be presented to volunteers in the project classification interface after all available data has been segmented.

### 4.7 Citizen science data export

The EAC project workflow examined in this manuscript was deactivated on 1st August 2019 and the project data exported from the data exports page of the Zooniverse Project Builder as a comma separated value (CSV) file. A single classification could consist of an arbitrary number of individual lines (a line being defined as a single continuous stroke annotation). Each segmentation is recorded in a JavaScript Object Notation (JSON) formatted string within the annotation field of the CSV file. Each line within the annotation is represented as a series of (x,y) pairs defining a line path (Supplementary Fig. 1F). The unaggregated segmentation data has been made available at www.ebi.ac.uk/biostudies/files/S-BSST448/etch-a-cell-classifications.csv.

### 4.8 Data aggregation with Contour Regression by Interior Averages (CRIA)

Multiple volunteer segmentations were produced for each slice uploaded to the EAC project. It was therefore necessary to remove outlying data and establish a ‘consensus’ segmentation for each slice. The Contour Regression by Interior Averages (CRIA) algorithm was developed to aggregate the volunteer segmentations. In this approach, first, each individual volunteer segmentation was converted into a closed loop. This procedure was performed for all segmentations associated with each slice of the ROI. Next, these closed loops were converted to interior areas and stacked. A consensus segmentation was determined by taking the outline of all interior areas where half or more of the volunteer segmentations were in agreement (Fig. 3A-F). This procedure was repeated for every slice within each ROI. Aggregated volunteer data has been made available at www.ebi.ac.uk/biostudies/files/S-BSST448/Aggregations. CRIA code is available at www.github.com/FrancisCrickInstitute/Etch-a-Cell-Nuclear-Envelope.

### 4.9 Model architecture

A U-Net CNN architecture [19, 20] was trained with aggregated volunteer segmentations for the automatic segmentation of the NE. This architecture uses convolutional layers and an autoencoder-style compression path. Hyper-parameter optimization was performed through random search, resulting in selection of the following model parameters; patch size of (12, 256, 256), dropout rate of 0.3, 32 start filters, Adam optimizer with an initial learning rate of 0.0005, and batch normalization. Informed by the expected and visible width of the membrane in the data (60-80nm), a nuclear envelope width of 70nm was selected.

Training data for the model consisted of the aggregated volunteer segmentations. A voxel resolution of 50nm was selected. Supplementary Table. 1 details the ROIs used for model training, validation and testing. Two ROIs were selected for model validation: Cell ID = C005 (ROI 2052-5784-112) and Cell ID = C010 (ROI 3588-3972-1). Two ROIs were used for model testing: Cell ID = C001 (ROI 1656-6756-329) and Cell ID = C006 (ROI 3624-2712-201); these two ROIs had been manually segmented by both expert and volunteers. One ROI (Cell ID = C013, ROI 1716-7800-517) was excluded from training due to having received insufficient data from the citizen science to perform data aggregation. Using a local high performance compute cluster (Methods, Nvidia Tesla V100-SXM2-32GB) to train this model took approximately 4 hours (1 hour for data pre-processing; 100 seconds per epoch = 3 hours in total).

The loss function used for the model was the smoothed dice coefficient (or F-measure), where:

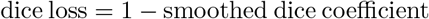

### 4.10 Model performance Metrics

A commonly applied approach to assess the quality of a model prediction versus ground truth for image data is to directly map the pixels between the two images. We report the F-measure, which is similar to the Dice coefficient. This measures the coincidence of predicted cell membrane to ground truth membrane. The F-measure of the nucleus area (as opposed to the NE) is also reported to enable easier comparison with previous work. As this metric requires a closed area, this was performed on a qualified single slice near the centre of the cell. Finally, we also report the Average Hausdorff Distance (AHD), in both pixels and nm, between the predicted NE and the position of the ground truth NE. This metric takes an average of all minimal distances between pixels in the prediction (P) and ground truth (G):

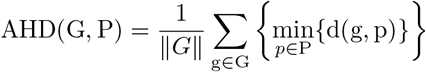

### 4.11 Tri-Axis Prediction (TAP)

A multi-axis modification of our machine learning model was implemented to improve performance (Supplementary Fig. 4). Data were downscaled in the xy plane to 50nm to be isotropic, and the stack transposed to run the model over each axis (Supplementary Movie. 6 and Supplementary Movie. 7). The resulting three orthogonal NE predictions were recombined to generate a final segmentation. All pixels assigned to NE in all three predictions were accepted, and connected components analysis was to remove over-segmented pixels (Methods). This approach was named ‘Tri-Axis Prediction’ (TAP). TAP results for each ROI have been made available at www.ebi.ac.uk/biostudies/files/S-BSST448/Predictions. TAP code is available at www.github.com/FrancisCrickInstitute/Etch-a-Cell-Nuclear-Envelope.

### 4.12 Post-processing of 3D volumes

Connected components analysis [35] was implemented as a post-processing step at multiple points within the analyses presented. Within the single-axis implementation of the machine learning model, connected components analysis was used to remove objects below a threshold of 10000 voxels. This threshold was selected as it resulted in the removal of small areas of erroneous over-segmented pixels, while legitimate membrane was preserved due to its comparatively large size. Connected components analysis was also used to isolate the predicted NE segmentation for the target nuclei within each ROI, to discard any predicted NE associated with peripheral cells potentially present. In the TAP modification of the machine learning model, connected components analysis was similarly used to remove over-segmented pixels by removing objects below a threshold of 10000 voxels and to identify and isolate the target nuclei within each ROI by selecting the largest connected component. The Python package, scikit-image ([36]) was used to automate these aspects of the data analysis pipeline (www.github.com/FrancisCrickInstitute/Etch-a-Cell-Nuclear-Envelope). Where NE segmentations produced by TAP are presented for the whole volume (e.g. Fig. 4J) objects below a more stringent threshold of 100000 voxels were removed using MorphoLibJ ([23]). 3D renderings of segmentations were generated using Fiji’s 3D Viewer plugin [37].

### 4.13 Computing Resource

Data analysis was performed on available high performance computing. This included Google Co-laboratory (www.colab.research.google.com), Amazon Web Services (AWS) cloud computing service (www.aws.amazon.com), and a local high performance compute cluster called “CAMP” (Crick Data Analysis and Management Platform). For reproducibility and convenience, the final analytical pipeline was packaged and tested on AWS.

### 4.14 Code availability

All assets relating to the analysis and training have been made available on public repositories and a single automated pipeline for reproducing the work has been containerised using Docker to capture environment configurations. The agnosticity of the containerised pipeline has been tested by running on a public cloud instance (AWS). Further information regarding re-running the pipeline has been provided in the readme on GitHub. We provide both reproducibility instructions (using the original data) and instructions for applying the trained model to other data sets. Code is available at: www.github.com/FrancisCrickInstitute/Etch-a-Cell-Nuclear-Envelope.

## Supporting information

SuppMovie_01

SuppMovie_02

SuppMovie_03

SuppMovie_04

SuppMovie_05

SuppMovie_06

SuppMovie_07

SuppMovie_08

SuppMovie_09

SuppMovie_10

SuppMovie_11

SuppMovie_12

SuppMovie_13

## 4.15 Author information

H.Sp., L.C. and M.J. conceived and directed the project. H.Sp. wrote the manuscript, with input from all authors. H.So., L.N., J.F., A.S., H.Sp and M.J. contributed data analysis. R.H., C.L. and H.Sp. contributed to EAC project development. C.J.P. and A.W. produced and analysed volume EM data. The Zooniverse volunteers contributed many segmentations, without which this project wouldn’t have been possible.

## 4.16 Acknowledgements

This work was supported by the Francis Crick Institute which receives its core funding from Cancer Research UK (FC001999), the UK Medical Research Council (FC001999), and the Wellcome Trust (FC001999). This publication uses data generated via the Zooniverse.org platform, development of which is funded by generous support, including a Global Impact Award from Google, and by a grant from the Alfred P. Sloan Foundation.

## 7 Supplementary Tables

**Supplementary Table 1:** Summary of volunteer effort applied to the 18 cells segmented in EAC. A total of 18 cells were manually identified for NE segmentation from the SBF SEM data (Fig. 1 and Supplementary Fig. 1).’Cell id’; unique cell identifier. ‘ROI’; region of interest ID. ‘No. subjects’; total number of slices for the cell. ‘Complete in z-axis’; describes whether the cell was complete along the z-axis. ‘Expert data available’; denotes whether an expert segmented the cell. ‘Total classifications’; the number of classifications that were made by volunteers for this cell. ‘Median classifications per subject’; the median number of classifications per subject. ‘Median classification time’; the median time spent by a volunteer on a classification for this cell (in seconds). ‘No. unique volunteers’; the number of unique volunteers contributing to the segmentation of the cell. ‘Total Time dd hh:mm:ss’; total volunteer time contributed. Only classifications taking between 0 and 2 hours were included when calculating timings. ‘Total time’; the amount of time spent by volunteers classifying the cell (in hours). ‘Machine learning’; denotes whether the segmentation data associated with the cell were used for training, validation or testing

| n | Cell id | ROI | No. subjects | Complete in Z axis | Expert data available | Total classifications | Median classifications per subject | Median classification time (seconds) | No. unique volunteers | Total Time dd hh:mm:ss | Total Time (hours) | Machine learning |
| --- | --- | --- | --- | --- | --- | --- | --- | --- | --- | --- | --- | --- |
| 1 | C001 | ROI_1656-6756-329 | 300 | yes | yes | 9300 | 39 | 61 | 2528 | 18 11:20:3 | 443 | Testing |
| 2 | C002 | ROI_1536-3456-213 | 300 | no | no | 8686 | 30 | 97 | 2741 | 21 23:4:16 | 527 | Training |
| 3 | C003 | ROI_2820-6780-468 | 199 | no | no | 4260 | 30 | 70 | 1380 | 8 15:50:39 | 208 | Training |
| 4 | C004 | ROI_2448-4704-271 | 300 | yes | no | 6096 | 30 | 81 | 1716 | 16 13:23:54 | 397 | Training |
| 5 | C005 | ROI_2052-5784-112 | 262 | no | no | 6258 | 30 | 84 | 1884 | 14 17:46:32 | 354 | Validation |
| 6 | C006 | ROI_3624-2712-201 | 300 | yes | yes | 8254 | 30 | 59 | 1515 | 15 7:11:35 | 367 | Testing |
| 7 | C007 | ROI_3576-5232-35 | 185 | no | no | 4669 | 30 | 50 | 1180 | 7 5:34:16 | 174 | Training |
| 8 | C008 | ROI_3516-5712-314 | 300 | yes | no | 7852 | 30 | 65 | 1694 | 16 6:4:23 | 390 | Training |
| 9 | C009 | ROI_1584-6996-1 | 151 | no | no | 3112 | 30 | 72 | 1283 | 6 14:12:3 | 158 | Training |
| 10 | C010 | ROI_3588-3972-1 | 151 | no | no | 2824 | 30 | 56 | 883 | 4 11:15:29 | 107 | Validation |
| 11 | C011 | ROI_3000-3264-393 | 274 | yes | no | 7377 | 30 | 56 | 1736 | 12 20:26:54 | 308 | Training |
| 12 | C012 | ROI_1608-912-1 | 151 | no | no | 3938 | 30 | 70 | 1569 | 8 9:3:16 | 201 | Training |
| 13 | C013 | ROI_1716-7800-517 | 150 | no | no | 233 | 2 | 101 | 110 | 0 11:45:53 | 12 | Not used |
| 14 | C014 | ROI_2832-1692-1 | 151 | no | no | 2010 | 30 | 62 | 817 | 3 17:23:53 | 89 | Training |
| 15 | C015 | ROI_1416-1932-171 | 300 | yes | no | 8173 | 30 | 98 | 2836 | 20 12:53:16 | 493 | Training |
| 16 | C016 | ROI_3768-7248-143 | 293 | no | no | 9243 | 30 | 53 | 1721 | 15 12:0:10 | 372 | Training |
| 17 | C017 | ROI_3972-1956-438 | 229 | no | no | 5434 | 30 | 62 | 1219 | 9 22:3:16 | 238 | Training |
| 18 | C018 | ROI_4320-1260-95 | 245 | no | no | 6869 | 30 | 66 | 1290 | 12 10:43:10 | 299 | Training |
| Total |  |  | 4241 |  |  | 104588 |  |  |  |  | 5137 |  |

## 8 Supplementary Figures

**Supplementary Figure 1:**
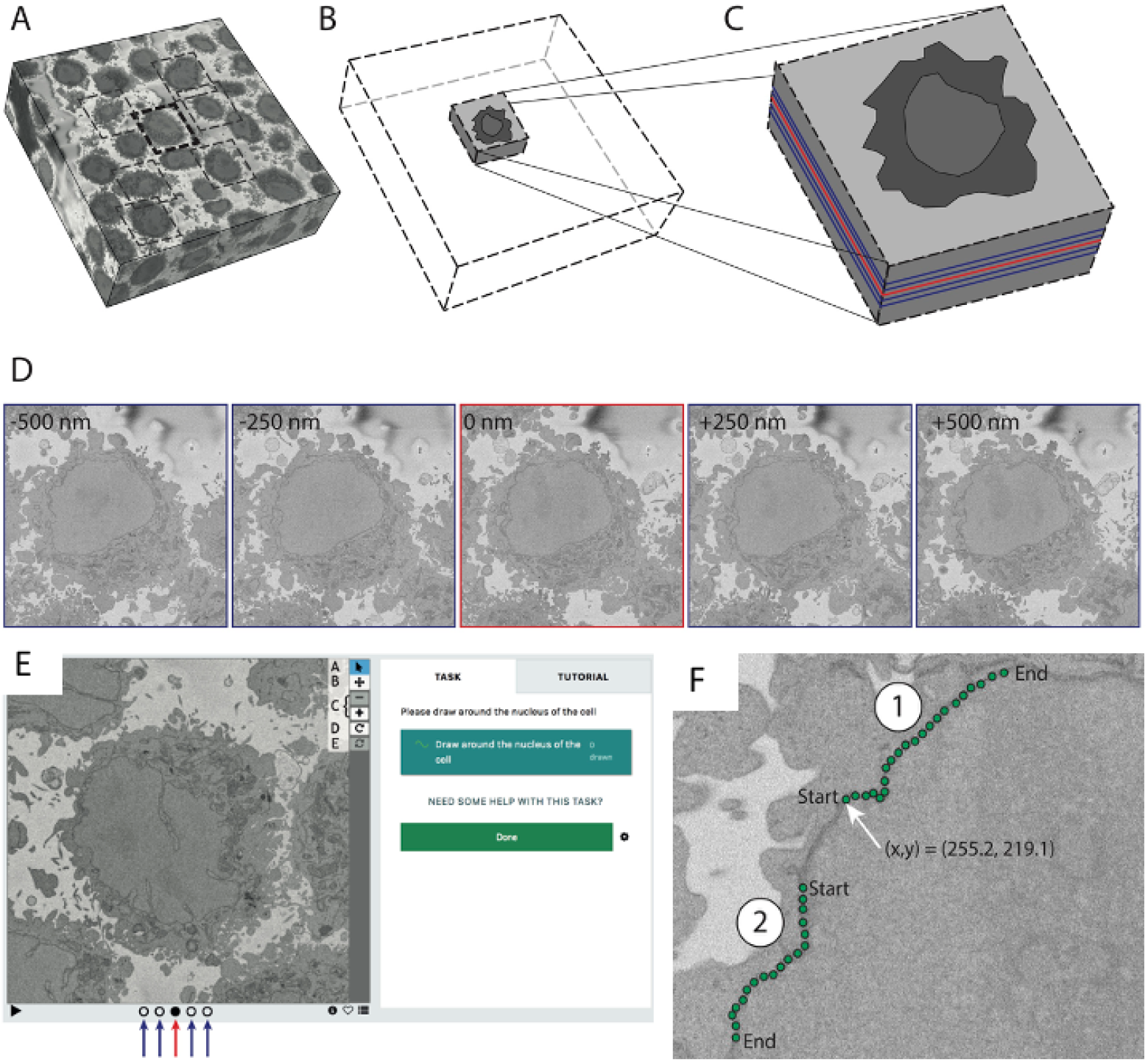
Data pre-processing and presentation to volunteers. Schematic of the process for subdividing the full raw data volume into cropped regions of interest (A); the raw data volume (8192 × 8192 × 518 voxels) is manually separated into regions of 2000 × 2000 pixels (dashed boxes). Each subvolume (B) is treated as a separate ROI (note, different ROIs can intersect in 3D). (C) Each ROI is a stack of up to 300 slices. (D) To provide 3D context to help resolve segmentation ambiguities, images are presented to volunteers as a flipbook containing 5 images; the central image (red) to be annotated along with slices 250nm and 500nm above and below (blue). The project classification interface (E) presents the volunteer with an image to be segmented at random from the selection of unretired images available. The image to be segmented (as indicated here with a red arrow) is shown as the central panel in a ‘flipbook’ of five images. (F) Schematic representation of the construction of the line vector objects; the green points illustrate the recorded (x,y) positions for a pair of separate path objects. A single user annotation may contain several separate lines which may or may not form a complete closed loop. The co-ordinate data are stored as separate x and y arrays for each separate path in a json formatted string within the values field of the CSV file.

**Supplementary Figure 2:**
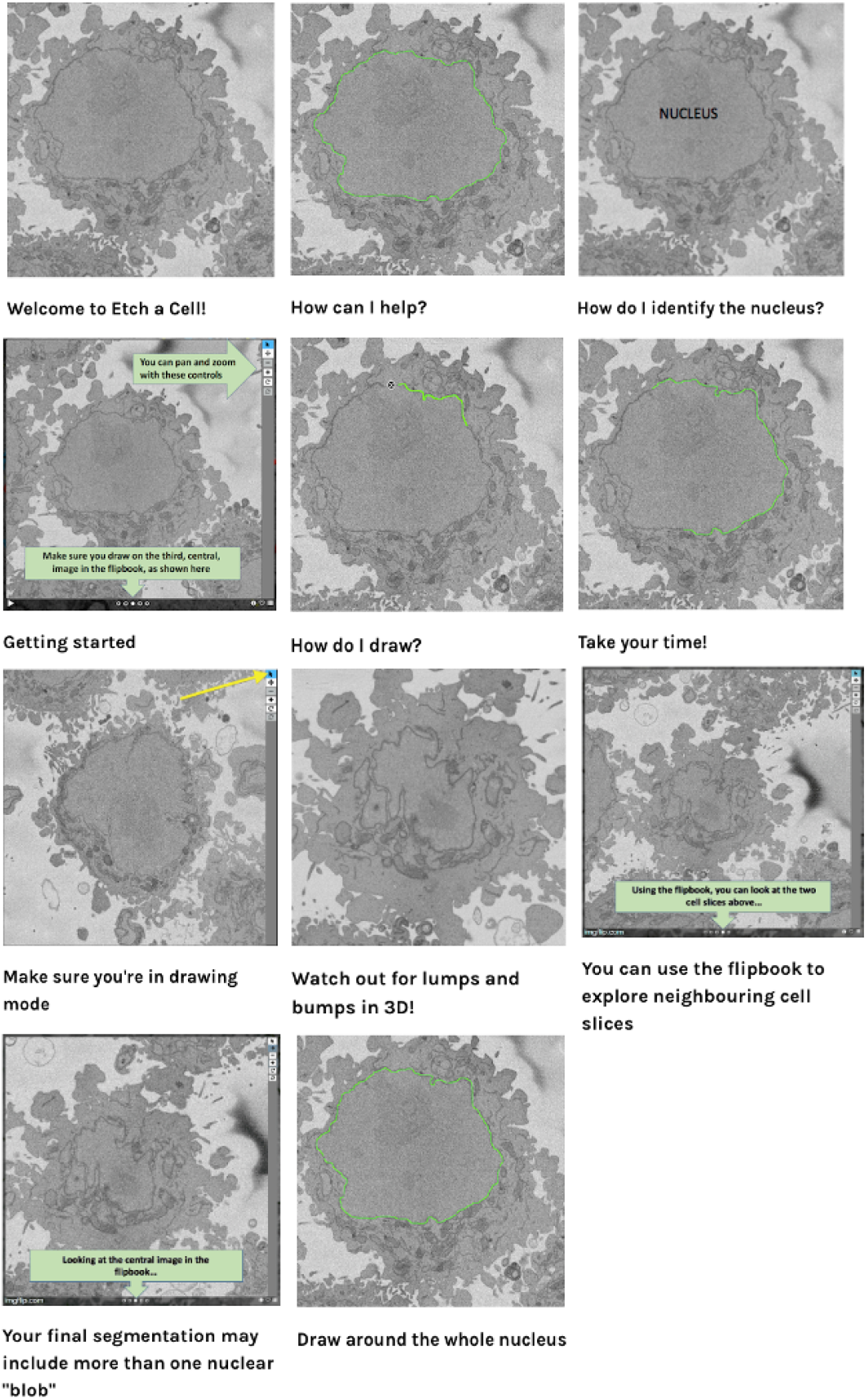
EAC project tutorial. Upon entering the classification interface of EAC for the first time, a volunteer is automatically presented with the project tutorial. Due to the complicated nature of NE segmentation task, the tutorial included ten comprehensive steps.

**Supplementary Figure 3:**
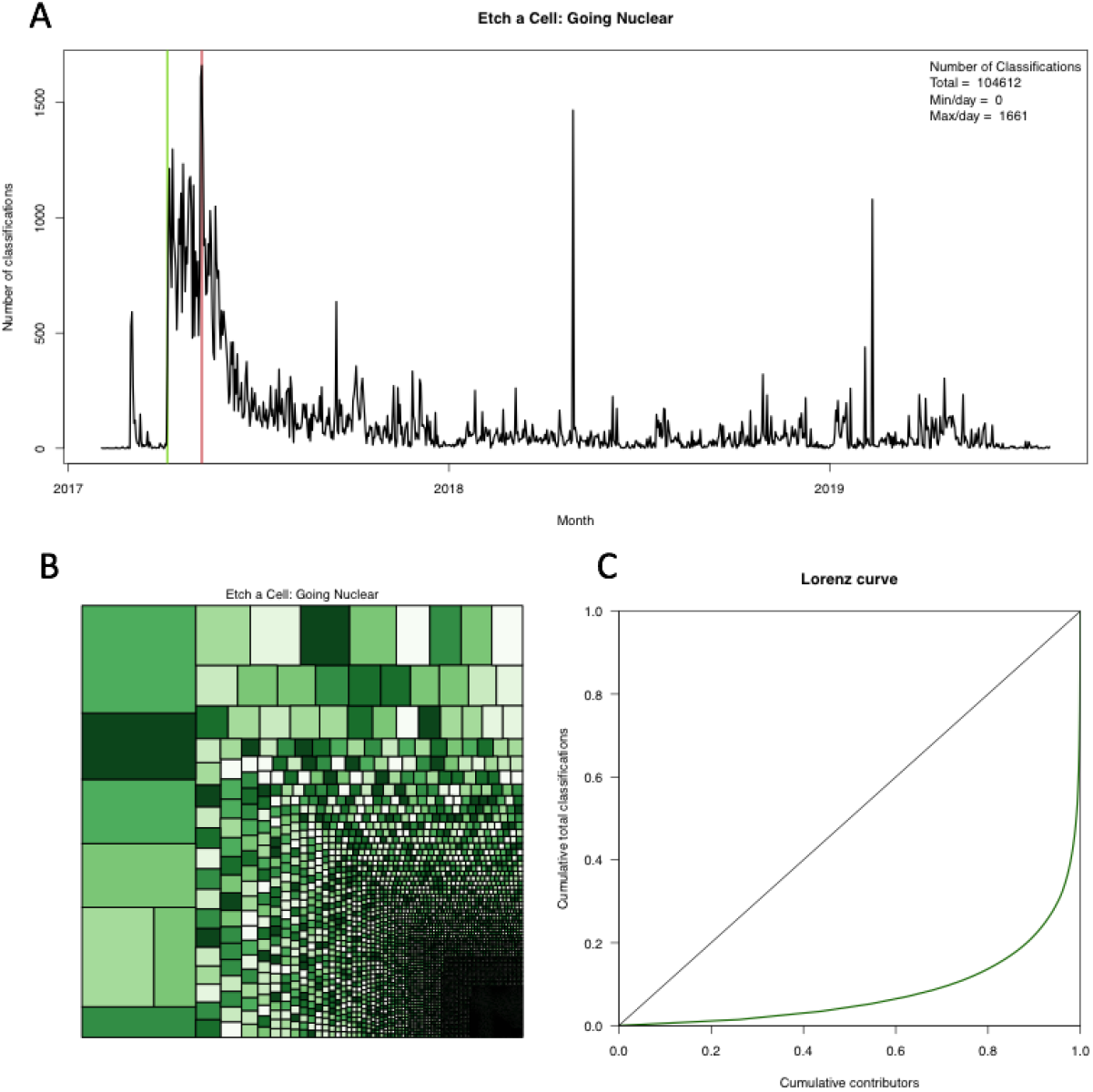
Volunteer interaction with EAC (A) Plotting classification count per day reveals a characteristic classification curve, with the majority of volunteer effort occurring shortly after project launch (indicated here with a green vertical line, 6th April 2017). The red vertical line indicates the day when most classifications were received, n = 1661, 9th May 2017). (B) A total of n = 4749 signed-in volunteers together contributed classifications to EAC. A large degree of variation was observed in the number of classifications submitted by each volunteer, as illustrated with this Treemap where the area of each cell corresponds to the total classifications submitted by a signed-in volunteer. (C) A Lorenz curve was plotted to describe the inequality in number of classifications per registered volunteer. This plot shows the cumulative number of classifications versus the cumulative number of volunteers, with the increased curvature of the Lorenz curve indicating stronger inequality in volunteer contribution. The black 45° line corresponds to total equality, which in this case would represent all users contributing equal numbers of classifications. The large distance here between the line of equality and the Lorenz curve illustrates the large amount of inequality between volunteers in the number of classifications made on EAC.

**Supplementary Figure 4:**
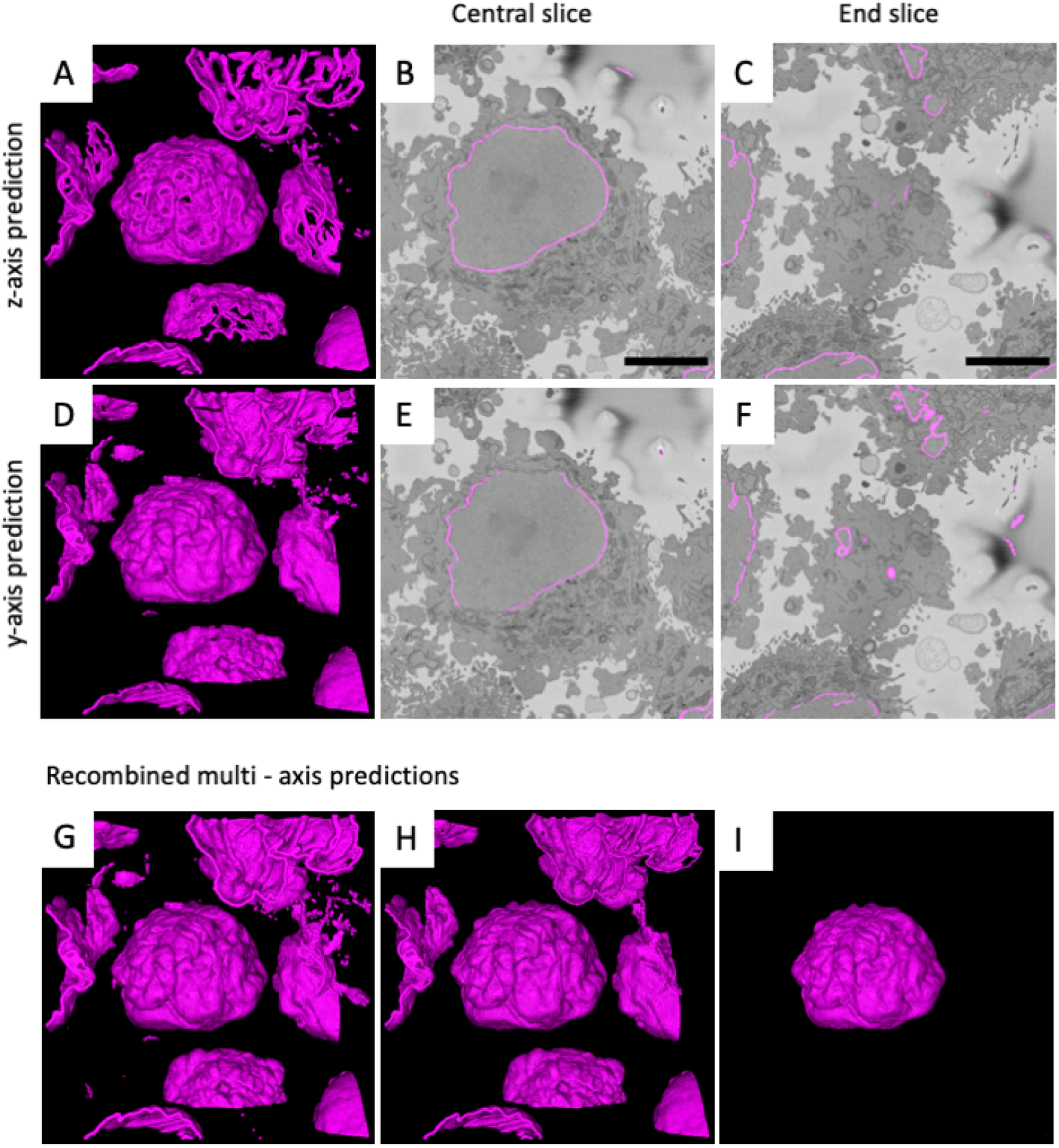
Tri-Axis Prediction for improved NE segmentation. Visualization of the predicted NE segmentation (A) revealed some regions of under-segmentation. These were not distributed randomly through the volume but instead related to predictable features of the changing NE structure, with central slices (B) proving easier to automatically segment compared to slices at the tops and bottom of the volume where the NE becomes more visually ambiguous, with the presence of islands and membrane parallel to the cutting plane (C). Because the data has been resampled to be isotropic, it was possible to transpose the EM stack to allow the model to be applied to the data in different orientations. Presenting the model with data in different orientations, e.g. along the y-axis (D), resulted in segmentation of NE pixels missed by the model when run other orientations (A). To enable visual comparison of model predictions, the y-axis segmentation prediction was transposed to the z-axis orientation (D-F). It can be be seen that the y-axis prediction (E) fails to capture some of the pixels correctly designated in the z-axis prediction (B) – these gaps correspond to the ‘ends’ when the structure is viewed from the y orientation. Conversely, the y-axis prediction (F) performs better for the z-axis ‘end slice’ (C). Predicting the NE along all three axes (x, y, z) and accepting all segmented pixels as NE results in a high-quality segmentation (G), however, a large number of erroneous, over-segmented pixels are also present that can be removed by connected components analysis (H). Connected components can also be utilized to select the central NE within an ROI through selecting the largest connected region (I). (5 micron scale bar is shown on panels B and C).

## 9 Supplementary Movies

Supplementary Movie 1: Full raw data volume. HeLa cells were imaged at 10nm pixel resolution with SBF SEM. The image stack consisted of 518 slices (with 50nm section thickness) collected at 8192 × 8192 pixels (Methods and Fig. 1). This volume was cropped in to ROIs for downstream analysis (Supplementary Fig. 1A-D and Supplementary Table. 1). The raw data associated with this volume have been made available via the EMPIAR repository (deposition ID: 137, accession code: EMPIAR-10094).

Supplementary Movie 2: All volunteer segmentations associated with one slice from one ROI. Each slice within each ROI was segmented by multiple volunteers (Fig. 2A-C). Although some variation in quality can be observed, with a three identifiable classes of error; ‘graffiti’, ‘false positive segmentation’ and ‘false negative segmentation’ (Fig. 2D-F), the majority of volunteer segmentations are distributed on and around the NE. Movie is produced from slice number 70 from C001 (ROI 1656-6756-329).

Supplementary Movie 3: 3D reconstruction of aggregated volunteer segmentations for each ROI. Individual volunteer segmentations for each slice within each ROI were aggregated with Contour Regression by Interior Averages (CRIA) (Methods and Fig. 3). This resulted in a high-quality segmentation produced by the collective effort of volunteers. Corresponding cell IDs can be seen in Fig. 3G.

Supplementary Movie 4: 3D reconstruction of expert, volunteer and machine predicted NE segmentations. Visual comparison of the 3D reconstruction of the NE for C001 (ROI 1656-6756-329) for expert, aggregated volunteer and machine predicted segmentations reveals a high similarity in NE segmentation (Fig. 4G-I).

Supplementary Movie 5: Expert, volunteer and machine predicted NE segmentations shown by slice for a single ROI. Visual inspection of expert, volunteer and machine-predicted NE segmentation per slice within C001 (ROI 1656-6756-329) reveals a high degree of overlap with the NE. Variation in segmentation quality can be observed in relation to the position of the slice within the stack, with slices found at the top and bottom of the volume displacing greater variation in segmentation quality (Fig. 4A-F). This is likely due to the presence of NE islands and membrane parallel to the cutting plane in these regions, which make these regions more challenging to segment.

Supplementary Movie 6: 3D reconstruction of TAP prediction for a single ROI. The data examined here were downscaled in the xy plane to 50nm to be isotropic, this enabled the stack to be transposed and the model to be run on each axis. The resulting three orthogonal NE predictions were recombined to generate a final segmentation. All pixels assigned to NE in all three predictions were accepted (Supplementary Fig. 4). This movie shows the recombined orthogonal predictions for a single ROI (C001). The NE pixels segmented by each of the three predictions are coloured coded as follows; x-axis = red, y-axis = green, and the z-axis = blue. Yellow regions are generated by a combination of red and green, and therefore indicate regions where the z-axis failed to segment NE pixels. Magenta regions are produced by red and blue, and therefore indicate where the y-axis has failed to segment NE pixels. Finally, cyan is generated from green and blue, therefore these regions indicate where the x-axis has failed to segment NE pixels (Supplementary Movie. 7).

Supplementary Movie 7: TAP prediction for a single ROI shown by slice. X, y and z-axis predictions for C001 are shown separately and in combination. The NE pixels segmented by each of the three predictions are coloured coded as follows; x-axis = red, y-axis = green, and the z-axis = blue. Therefore, in the combined panel, yellow regions are generated by a combination of red and green, and therefore indicate regions where the z-axis failed to segment NE pixels. Magenta regions are produced by red and blue, and therefore indicate where the y-axis has failed to segment NE pixels. Finally, cyan is generated from green and blue, therefore these regions indicate where the x-axis has failed to segment NE pixels (Supplementary Movie. 6).

Supplementary Movie 8: 3D reconstruction of the predicted NE segmentation for the full data volume. Automatic NE segmentation with TAP (Methods) identified many nuclei which had not previously been segmented through expert or volunteer effort (Fig. 4J), including a mitotic cell Fig. 5A).

Supplementary Movie 9: Predicted NE segmentation for the full data volume shown by slice. The application of TAP to the full data volume identified many nuclei which had not previously been segmented through expert or volunteer effort (Fig. 4J).

Supplementary Movie 10: 3D reconstruction of the predicted NE segmentations for each ROI. The NE of each ROI was automatically segmented using TAP (Methods). Corresponding cell IDs can be seen in Fig. 4K.

Supplementary Movie 11: Predicted NE segmentation for a mitotic cell within the full data volume shown by slice. Applying TAP to the full data volume allowed us to observed that the model performed well when segmenting the partially broken down NE of a mitotic cell (Fig. 5A).

Supplementary Movie 12: Predicted NE segmentation for 5nm SBF SEM data shown by slice. TAP was applied to an alternative region from the same resin-embedded sample imaged at higher resolution (5nm) on the same microscope (Fig. 5B). Visual inspection showed good model performance, indicating the model is generalisable to novel contexts. However, some erroneous over-segmented pixels can be observed in this data, indicating there is room for future improvement.

Supplementary Movie 13: Predicted NE segmentation for HeLa cells imaged with FIB SEM data, shown by slice. TAP was applied to a HeLa cell from the same sample imaged by an alternative volume EM methodology (FIB SEM) (Fig. 5C). Visual inspection showed good model performance.

